# Interregional causal influences of brain metabolic activity reveal the spread of aging effects during normal aging

**DOI:** 10.1101/490292

**Authors:** Xin Di, Marie Wölfer, Mario Amend, Hans Wehrl, Tudor M. Ionescu, Bernd J. Pichler, Bharat B. Biswal, the Alzheimer’s Disease Neuroimaging Initiative

## Abstract

During healthy brain aging, different brain regions show anatomical or functional declines at different rates, and some regions may show compensatory increases in functional activity. However, few studies have explored interregional influences of brain activity during the aging process. We proposed a causality analysis framework combining high dimensionality independent component analysis (ICA), Granger causality, and LASSO (least absolute shrinkage and selection operator) regression on longitudinal brain metabolic activity data measured by Fludeoxyglucose positron emission tomography (FDG-PET). We analyzed FDG-PET images from healthy old subjects, who were scanned for at least five sessions with an averaged intersession interval of about one year. The longitudinal data were concatenated across subjects to form a time series, and the first order autoregressive model was used to measure interregional causality among the independent sources of metabolic activity identified using ICA. Several independent sources with reduced metabolic activity in aging, including the anterior temporal lobe and orbital frontal cortex, demonstrated causal influences over many widespread brain regions. On the other hand, the influenced regions were more distributed, and had smaller age related declines or even relatively increased metabolic activity. The current data demonstrated interregional spreads of aging on metabolic activity at the scale of a year, and have identified key brain regions in the aging process that have strong influences over other regions.

## 1. Introduction

The human brain undergoes development and aging across the entire life-span. Neuroimaging studies have demonstrated that different brain regions develop and age in different rates. The global gray matter volume decreases linearly after 20s of age, but some regions such as the bilateral insula, superior parietal gyri, central sulci, and cingulate sulci show faster volumetric declines as measured by voxel-based morphometry (Good et al., 2001). Cortical thickness measures show more widespread cortical thinning patterns during aging (Salat et al., 2004). In contrast, results from functional MRI studies (fMRI) are more complex with some brain regions show increased activations in certain tasks rather than declined activations accompanied with the anatomical declines (Di, Rypma, & Biswal, 2014; Spreng, Wojtowicz, & Grady, 2010). In addition, the functional alterations in aging may depend on the task domains and behavioral performances (Spreng et al., 2010), making it difficult to conclude a region to be functionally increased or decreased in aging. A complementary approach is to study brain activity during a state without specific behavioral involvements, i.e. resting-state. Studies have been performed earlier using positron emission tomography (PET) (Kuhl, Metter, Riege, & Phelps, 1982; Martin, Friston, Colebatch, & Frackowiak, 1991; Zuendorf, Kerrouche, Herholz, & Baron, 2003), and later using resting-state fMRI (Biswal et al., 2010). Using a large sample of over 1,000 subjects, Biswal and colleagues have showed reduced resting-state activity in aging mainly in the default model network and increased activity in the visual, motor, and subcortical regions (Biswal et al., 2010).

Functionally related brain regions typically show similar co-developments (Alexander-Bloch, Raznahan, Bullmore, & Giedd, 2013) or co-declines, therefore yielding cross-subject interregional covariances. This has been shown as early as 1980s using regional cerebral blood flow data (Prohovnik, Håkansson, & Risberg, 1980) and regional metabolic activity data (Horwitz, Duara, & Rapoport, 1984; Metter, Riege, Kuhl, & Phelps, 1984) measured using PET. Later, more sophisticated methods, such as independent component analysis (ICA) and graph theory based analysis, have been applied to PET data to study the brain metabolic covariance networks (Di et al., 2017; Di & Biswal, and Alzheimer’s Disease Neu, 2012). Correlated metabolic activity or blood flow was typically found between left/right homotopic regions, and between some within hemisphere regions that are functionally related, e.g. language related regions in the left hemisphere. However, different connectivity patterns have been found between this metabolic covariance connectivity and the resting-state connectivity that has been typically observed from fMRI data (Di et al., 2017; Di & Biswal, and Alzheimer’s Disease Neu, 2012). The inter-subject covariance patterns have also been shown using other imaging modalities, such as brain volumes (Di & Biswal, 2016; Douaud et al., 2014; Mechelli, Friston, Frackowiak, & Price, 2005), cortical thickness (Lerch et al., 2006), and different resting-state fMRI indices (P. A. Taylor, Gohel, Di, Walter, & Biswal, 2012; Zhang et al., 2011).

Given the different rates of declines or relative preservations of different brain regions in aging, and large scale brain networks working in synchrony during both task execution and resting-state (Bullmore & Sporns, 2009, 2012; Di, Gohel, Kim, & Biswal, 2013), it is likely that the regions that are working together affect each other during the aging process. Specifically, a region that declines faster may influence another region during functional interactions in everyday basis; therefore would cause the other region to decline or show a compensatory increase of functional activity. So, it is critical to study the causal interregional influences between regions rather than the simple covariance, especially at the time scale of months to years when brain aging could be observed. Although regional brain aging is generally assumed to be linear in trend, the observed regional brain measures might showed fluctuations along the linear trend (Figure 1). The causal influence between regions could then be captured by causality analysis methods such as Granger causality (Granger, 1969). By using Granger causality we could examine whether the brain activity in a brain region at time points of months or years earlier can predict the activity of another brain region at the current time point. Granger causality at the similar time scales has been studied on brain morphological progressions in epilepsy (Zhang et al., 2017) and schizophrenia (Jiang et al., 2018) based on anatomical MRI data. However, both of these studies are cross-sectional. Large-scale multi-site longitudinal open access dataset, such as Alzheimer’s Disease 4 Neuroimaging Initiative (ADNI), has made it possible to examine causal influences during aging in a within-subject manner. Extending our previous work on metabolic covariance networks using Fludeoxyglucose (FDG) PET images (Di et al., 2017; Di & Biswal, and Alzheimer’s Disease Neu, 2012), we sought in the current study to examine the interregional causal influences of metabolic activity during aging.

**Figure 1.**
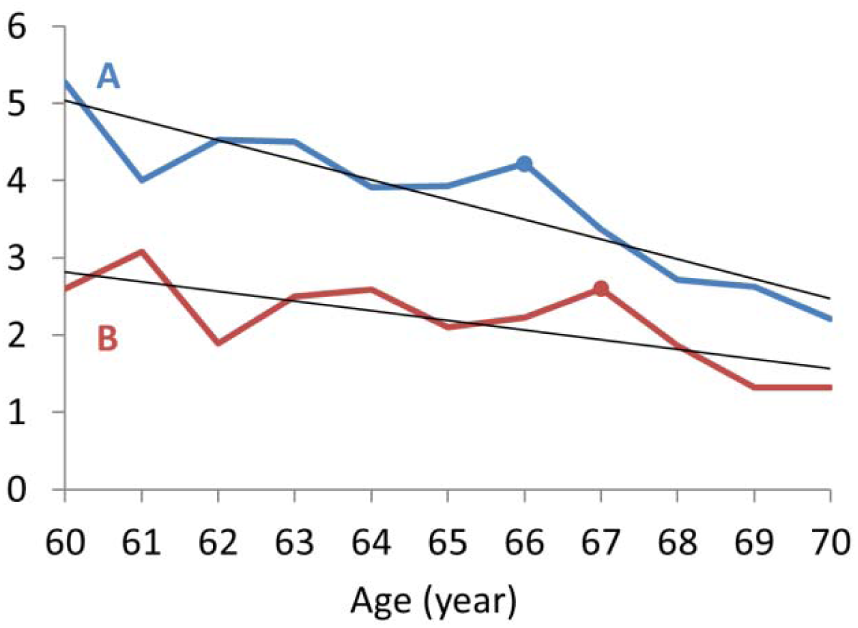
An illustration of interregional causal effects during aging. Regions A and B both show linear declines during aging at different rates, with additional fluctuations along the linear trends. The fluctuations of region A influences those in region B, so that an event in A can be observed one time point later in B (e.g. the marked peak at age 66 and 67 in A and B, respectively).

The aim of the current study is to explore the causal interregional influences of metabolic activity during normal aging at the time scale of a year. We leveraged the longitudinal FDG-PET data from the ADNI dataset, where there were at least five sessions of FDG-PET scans for each subject at a time step of approximately one year. First, we examined regional age effects of metabolic activity to identify regions with accelerated declines, with no apparent age effects, and with relative increases. Second, we performed whole brain Granger causality analysis to identify causal influences, where the metabolic activity in a region at a certain time point can be predicted by the metabolic activity in another region at the previous time point. We predict that the regions that show accelerated declines during aging will cause other regions to decline, thus showing interregional spreads of age effects.

## 2. Materials and methods

### 2.1 ADNI data

Data used in the preparation of this article were obtained from the ADNI database (ADNI - Alzheimer’s Disease Neuroimaging Initiative: RRID:SCR_003007; adni.loni.usc.edu). The ADNI was launched in 2003 as a public-private partnership, led by Principal Investigator Michael W. Weiner, MD. The primary goal of ADNI has been to test whether serial magnetic resonance imaging (MRI), positron emission tomography (PET), other biological markers, and clinical and neuropsychological assessment can be combined to measure the progression of mild cognitive impairment (MCI) and early Alzheimer’s disease (AD). For up-to-date information, see www.adni-info.org.

Only data from healthy participants were included in the current analysis. All participants showed no signs of depression, mild cognitive impairment, or dementia, with Mini-Mental State Exam (MMSE) scores between 24 and 30 and Clinical Dementia Rating (CDR) score of 0. We manually selected longitudinal FDG-PET images from the ADNI database, with participants who had at least five sessions of FDG-PET images available. As a result, 72 subjects (25 females) were included in the current analysis with a total of 432 PET scan sessions. The numbers of available sessions ranged from 5 to 9 (Figure 2A). The average age at the first session was 75.8 years (62 to 86 years). For each session, we calculated a mean image or adopted the only image to represent the session. The main analysis focused on the changes and causal influences over the time scale of sessions. The intersession interval with a subject varied from 3 months to up to 8 years for a few rare cases (Figure 2B). The mean and median of the intersession intervals were 1.02 and 0.98 years, respectively.

**Figure 2.**
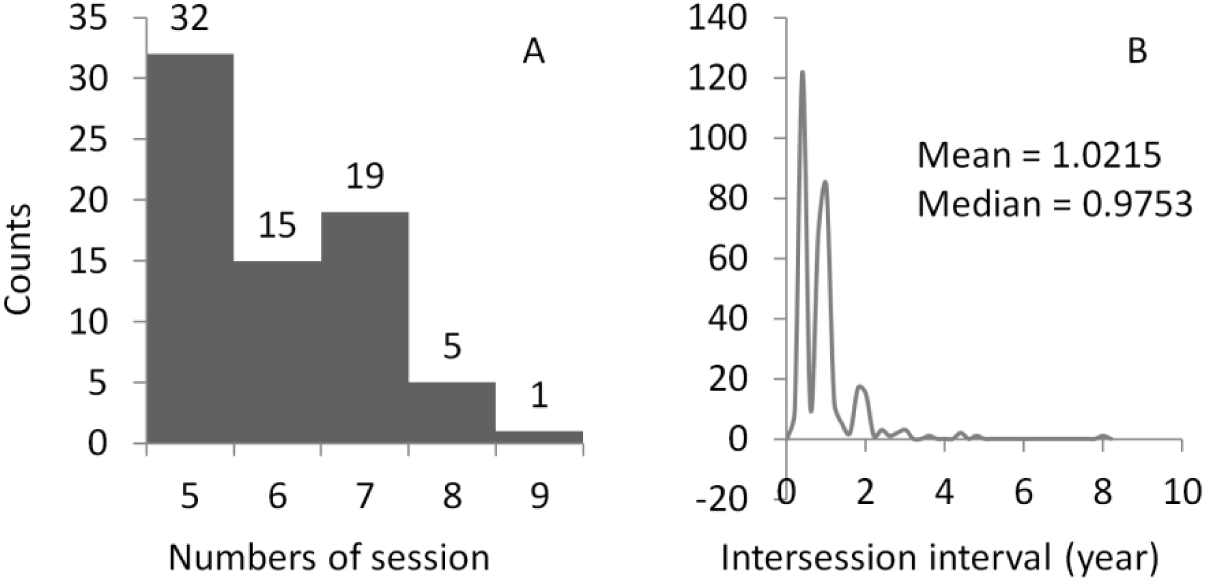
Histograms of the numbers of sessions for each subject (A) and the intersession intervals for all the sessions and subjects (B). The participants were typically studied at 0, 6, 12, 24, 36 months related to the first visit, and yearly follow-ups. Therefore, the intersession intervals are likely to be around six months or one year.

The FDG-PET images were acquired from multiple sites with different PET imaging protocols. However, the imaging parameters were mostly similar across different sessions within a subject. Since the current analyses were all within-subject, the impacts of different imaging parameters from different sites can be effectively minimized. More information about the PET protocol can be found in (Jagust et al., 2010). All the images and subjects included in the current analysis can be found at: https://osf.io/4a3vt/.

### 2.2. PET data preprocessing

The PET data were preprocessed using SPM12 (SPM: RRID:SCR_007037; https://www.fil.ion.ucl.ac.uk/spm/) under MATLAB R2017b. For each subject, if there were more than one PET image in a session, all the PET images in the session were realigned to the first image and the mean image of the session was calculated. The mean images (or the only image) across all the sessions of a subject were then realigned to the one in the first session. The cross-session mean image was normalized directly to the PET template in SPM in standard Montreal Neurological Institute (MNI) space, and then all the images were normalized to MNI space using the same set of parameters. We chose the direct normalization approach rather than using an anatomical MRI as a mediator, because the spatial resolutions of the PET images were adequate and the direct normalization has its own advantage compared with the anatomical MRI mediated method (Vince D. Calhoun et al., 2017). The images were then spatially smoothed using a Gaussian kernel with 8 mm FWHM (full width at half maximum). Lastly, each image was divided by its mean signal within an intracranial volume mask.

### 2.3. Independent component analysis

We first performed spatial ICA to separate the whole brain metabolic maps into independent sources of local metabolic variations (Di & Biswal, and Alzheimer’s Disease Neu, 2012). We extracted a relatively high number of ICs, so that the resulting ICs could represent more local variations than large scale networks (Fu et al., 2018, 2019; Smith et al., 2013). This data-driven approach is an alternative to atlas-based parcellation, and may be more representative to local variations of metabolic activity. The ICA was performed using Group ICA of fMRI Toolbox (GIFT: RRID:SCR_001953; http://mialab.mrn.org/software/gift) (V D Calhoun, Adali, Pearlson, & Pekar, 2001). The preprocessed FDG-PET images from different sessions and subjects were concatenated into a single time series, and fed into the ICA analysis. Eighty one components were recommended by the minimum description length (MDL) algorithm implemented in GIFT. After IC extraction of the 81 components, the ICs were visually inspected and grouped into eight domains (Supplementary materials) as well as 21 noise components. There were in total 60 ICs included in the following analysis. For each IC, the associated time series were obtained to represent metabolic activity of this source in different subjects and sessions. The 81 IC maps are available at: https://osf.io/4a3vt/.

### 2.4. Regional age effects

For each subject, a general linear model (GLM) was built to examine the linear aging effects. The GLM included two regressors, a constant term and a linear age effect. The GLM analysis was performed on each IC, and the *β* values of the age effect were obtained. Group level analysis was then performed on each IC using a one sample t test model to examine the group averaged effect of age. A FDR (false discovery rate) corrected p < 0.05 was used to identify ICs that had significant age effects after correcting for all the 60 comparisons. Thereafter, the ICs were sorted into three groups, with significant (relative) increased metabolic activity, with no significant changes, and with significant decreased metabolic activity in aging.

### 2.5. Interregional causality analysis

We adopted Granger causality like method to examine the interregional causal influence of metabolic activity. Specifically, we treated the longitudinal FDG-PET data as time series, and used autoregressive model to predict the value of time point t in a region *y* by the previous time points of another region *x*, when controlling for its own previous time points. In the current data, the time step is approximately one year. To account for the variability of intersession interval, the intervals between time points t and t – 1 were added as a covariate or regressed out in the analysis (see below for details). Another consideration is the order of the model, i.e. how many previous time points are used to predict the current time point. In this study, we used only the first order model to measure the causal influence of only one previous time point, which represents a time step of about one year. The limited number of time points in a subject prevents us to use higher order models. The advantage of using only the first order model is that the sign information of the beta estimate enables us to differentiate positive and negative predictions. The model can be expressed in the following form:

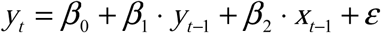

where *y* represents the predicted time series in one brain region, and *x* represents the predicting time series of another brain region. *y*_t-1_ represent the time series of *y*_t_ which moved one time point ahead, thus representing a autoregression model of time series *y*. The effect of interest is the predicting value of *x*_t-1_, which is β_2_.

We concatenated the time series across all the subjects to form a long time series for analysis (Figure 3). Therefore, the model is considered fixed effect model. The first to m_i_ time series of a subject were first z transformed to minimize inter-subject variation, where m_i_ represents the total number of time points in a subject i. For each subject, we included the time points 2 to m of the time series of a region as the predicted variable *y*_t_. The autoregressive variable *y*_t-1_ of the same region included the time points 1 to m − 1 of the time series of the same region. The predicting variable *x*_t-1_ was the time points 1 to m – 1 from another region *x*. After concatenation, there were in total 360 data points in the time series.

**Figure 3.**
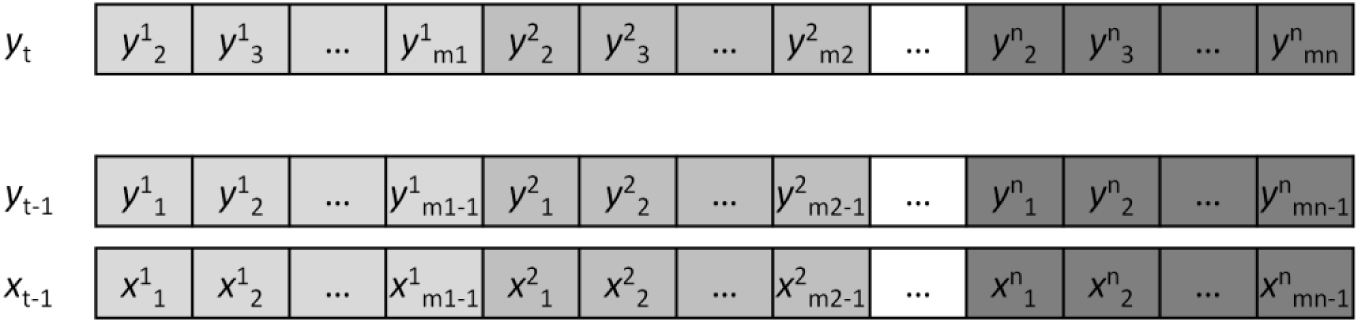
Illustration of the construction of the variables used in the causality analysis. X represents the predicting region, and y represents the predicted region. The superscript represents different subjects, with a total number of n. The subscript represents the scan session in a subject, with a total number of mi for a subject i.

The model could be applied to each pair of the ICs from the 60 ICs. The intersession interval between time *t* and time *t – 1* were included in the model as a covariate. We first performed such analysis on each pair of ICs to obtain the predicting effect (*β*_2_) and corresponding p values, which formed a 60 x 60 matrix of causal effects. FDR correction at p < 0.05 was used to correct for multiple comparisons of the in total 3,540 (60 × 59) effects, where autoregressive effects along the diagonal were not tested.

This pair-wise approach may identify influences from different regions with shared variance although maybe only one region has direct influence with the tested region. To overcome this, when predicting a region *x*^i^_t_, one can add all the other ICs to identify which region can predict *x*^i^_t_. The model is then as following for a predicted region *x*^i^:

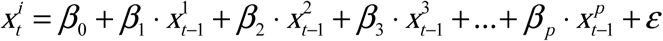

Since all the ICs were included in the model, there was no need to differentiate the variables of *x* and *y*. Therefore, we use *x* to denote all the time series variables. The superscripts of *x* now represent different ICs, where p represents the total number of the IC. Before entering in to the model, a time series representing the intersession interval between time point t and t – 1 were regressed out from all the *x*_t-1_ time series to account for the intersession interval variability. Estimating the multivariate model may be challenging, especially when some of the IC time series may be highly correlated. It can be assumed that only a small number of ICs may influence the predicted IC. In this scenario, one can use regularization method to estimate the sparse influence effects, such as using LASSO (least absolute shrinkage and selection operator) (Tang, Bressler, Sylvester, Shulman, & Corbetta, 2012; Tibshirani, 1996). The motivation of choosing LASSO over other regularization methods is that the LASSO regularization can force some parameters in the model to be zero thus resulting in only a small number of non-zero parameters. This is important in the current context, because the aim is to identify a small number of interregional influences. Since this model examines the prediction of the time series of one IC by the time series of all the other ICs, the analysis only needed to be performed for 60 times (compared with 60 x 59 times in the pair-wise analysis) to cover all the ICs.

The LASSO regression was performed using the lasso function implemented in MATLAB. To determine an optimal regularization factor *λ*, we used a set of *λ* from 0 (no regularization) to 0.5 with a step of 0.001. The identified non-zero influences dropped dramatically as the increase of *λ*. We identify the *λ* where the number of non-zero influences were the closest to the number of significant effects when using FDR correction in the pair-wise analysis, and reported all the non-zero influencing effects.

The resulting 60 × 60 influencing matrix can be treated as a directed network graph, where the ICs represent the nodes and the causal influences represent directed edges of the graph. We calculated in-degree and out-degree of the 60 nodes to characterize the importance of an IC in the whole brain influencing graph. To ensure that the degree calculation was not affected by arbitrary defined threshold, we also explored the graphs from other *λ* values to verify the identified hub regions are still present. To visualize the network topology, we identified the giant component where all the nodes in the component were somehow connected (without considering the direction of the influences). The giant components were visualized using the force layout.

## 3. Results

### 3.1. Age effect on regional metabolic activity

We first examined the age effects on the regional metabolic activity for the 60 ICs. Statistical significant ICs at p < 0.05 after FDR correction are shown in Figure 4. Eighteen ICs showed significant reduced metabolic activity, including one IC that covered the inferior portion of the cerebellum (IC 1), three ICs that coved visual cortex (IC 17, 32, and 53), three ICs that covered the posterior parietal cortex (IC 21, 27, and 28), five ICs that covered the anterior portion of the temporal lobe and insula (IC 24, 44, 63, 66, and 77), one IC that covered the thalamus and basal ganglia (IC 23), two ICs that covered the orbital frontal cortex and frontal pole (IC 3 and 48), and three ICs that covered the cingulate cortex and neighboring midline cortical regions (IC 7, 37, and 46). The left panel of Figure 5 illustrates the negative age effects of an example IC (IC 48). It can be seen that there is a general linear trend of decrease of metabolic activity. But each subject showed fluctuations of metabolic activity along the linear trend. In contrast, 6 ICs showed increased metabolic activity. It should be noted that due to the nature of PET imaging, the global signal for each PET image has to be normalized. Therefore, it is difficult to say whether the positive age effect represents increased metabolic activity, or a relative increase with reference to the global effect. The ICs with relative increased metabolic activity during aging included one IC covering the inferior and posterior portion of the cerebellum (IC 13), two subcortical ICs covering the basal ganglia, insula, amygdala, and thalamus (IC 19 and 52), and three ICs of sensorimotor regions (IC 36, 50, and 68). The right panel of Figure 5 illustrates the positive age effects of an example IC (IC 19). There were 36 ICs that did not show statistically significant age effects at p < 0.05 after FDR correction. The middle panel of Figure 5 illustrates the age effects of an example IC (IC 56) with no statistical significant age effect.

**Figure 4.**
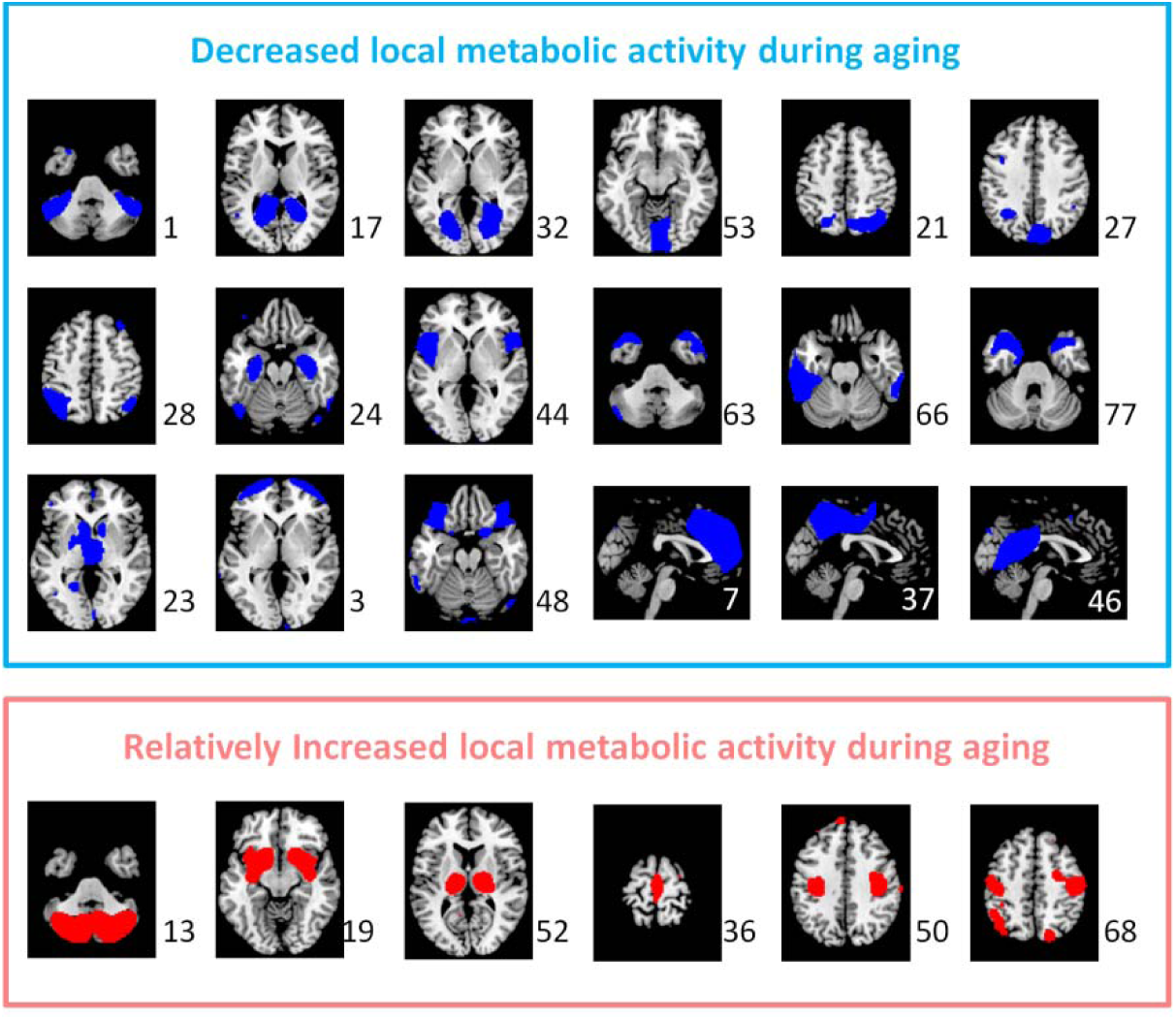
The independent components (ICs) that showed statistically significant decreased (blue) and increased (red) metabolic activity during aging after controlling for global effect at p < 0.05 with false discovery rate (FDR) correction. The numbers to the bottom right represent the IC number.

**Figure 5.**
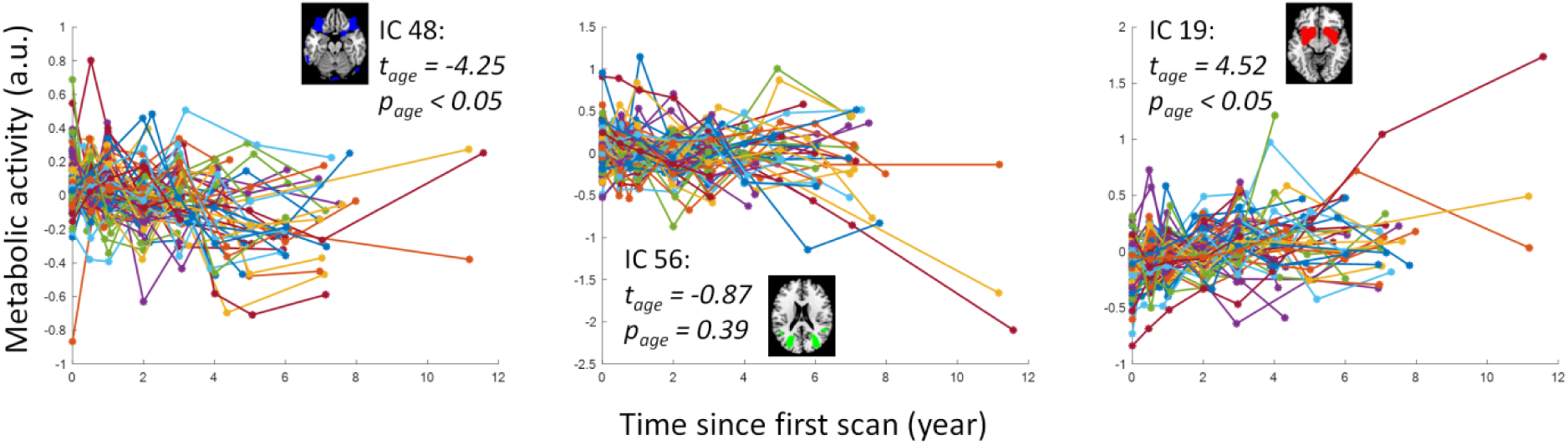
Examples of aging effects of metabolic activity of three independent components (ICs) that had negative, non-significant, and positive aging effects. Each colored line represents one subject. T and p values represent group-level one sample t test statistics. A.u., arbitrary unit.

### 3.2. Interregional causal influences of metabolic activity

We first applied pairwise autoregressive model to obtain a 60 × 60 matrix of the interregional causal influences of metabolic activity between each pair of the ICs (Figure 6A). When using a statistical threshold of p < 0.05 of FDR correction, 14 positive and 13 negative causal influences were identified (Figure 6B). We next performed LASSO regression with *x*_t_ of an IC as the predicted variable and *x*_t-1_ of all the ICs as the predicting variables using a range of *λ*. We identified the *λ* value where the number of non-zero effects was the closest to the number of significant effects in the pairwise analysis. The resulting influencing effects at *λ* = 0.162 (Figure 6C) look in general similar to the significant effects identified by the pairwise analysis, although some subtle differences can be noted. There were 15 positive and 13 negative causal influences identified using the LASSO regression method (Table 1).

**Figure 6.**
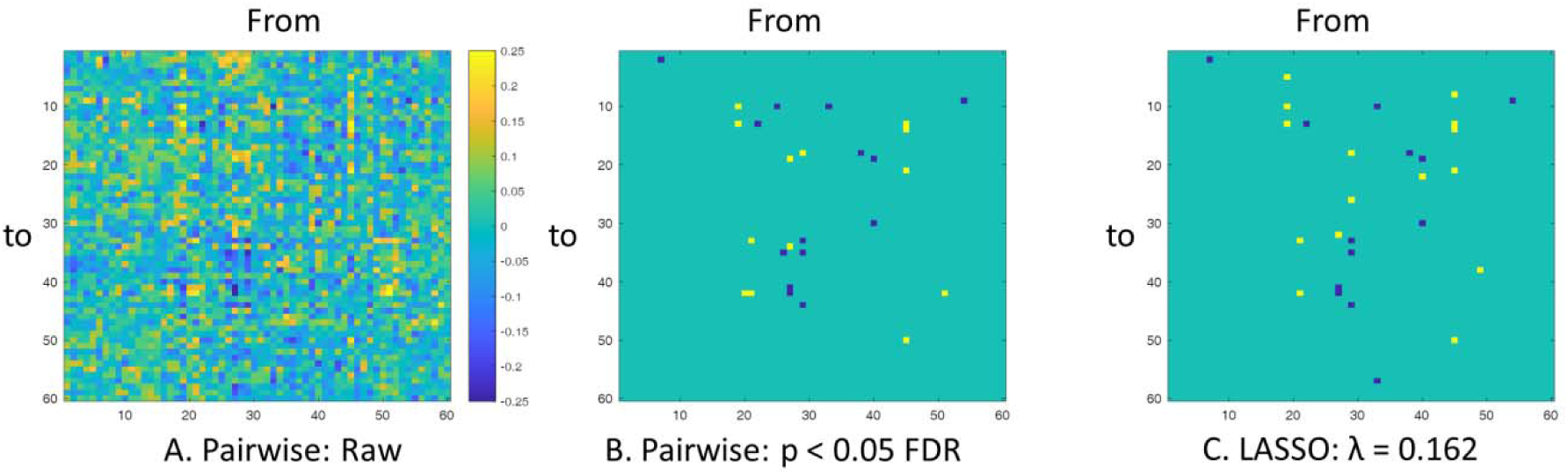
A, Pairwise matrix of interregional causal influence of metabolic activity. The columns represent influencing independent components (ICs), while the rows represent influenced ICs. B, Ternary matrix of significant positive or negative interregional causal influences thresholded at p < 0.05 after false discovery rate (FDR) correction. C, Ternary matrix of positive or negative interregional causal influences identified at *λ* = 0.162 using LASSO (least absolute shrinkage and selection operator) regression.

**Table 1.**
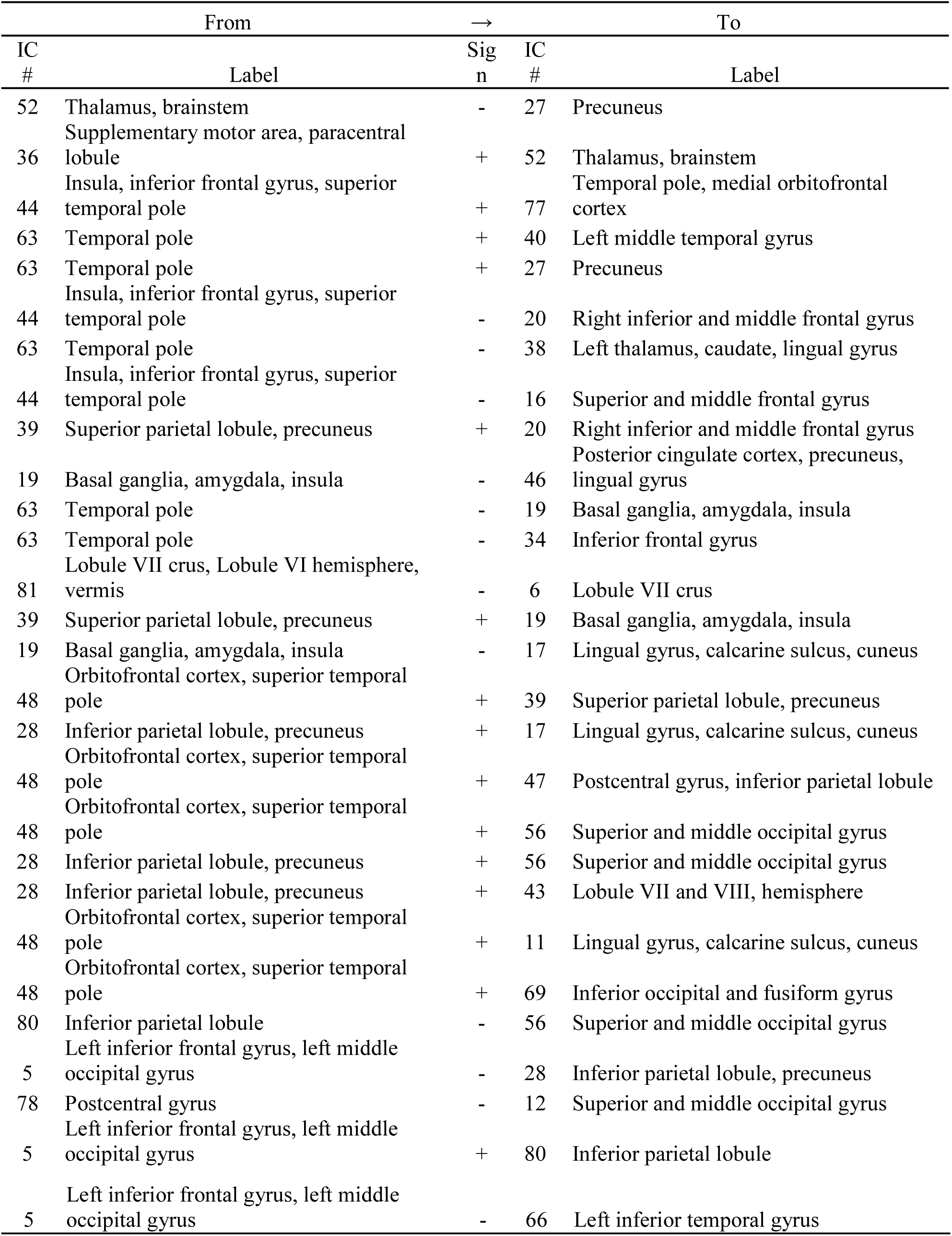
List of interregional causal influences of metabolic activity identified at λ = 0.162 using LASSO (least absolute shrinkage and selection operator) regression.

Among the 28 causal influences from the LASSO regression, the first giant component was comprised of 26 causal influences involving 25 ICs (Figure 7). The IC maps were color coded based on their regional age effects to illustrate the relationships between regional metabolic activity changes and the signs of causal influences. It can be seen that the influences between two decreased regions or two increased regions in aging were in general positive, but the influences between one increased and one decreased regions were in general negative. For example, the bilateral anterior temporal IC (IC# 63 in Figure 6) positively influenced the medial parietal IC (IC# 27), but negatively influenced the basal ganglia IC (IC# 19). It is consistent with the direction of the spread of age effects. There were also ICs that without apparent age effects, where the signs of causal influences with other regions did not show clear pattern.

**Figure 7.**
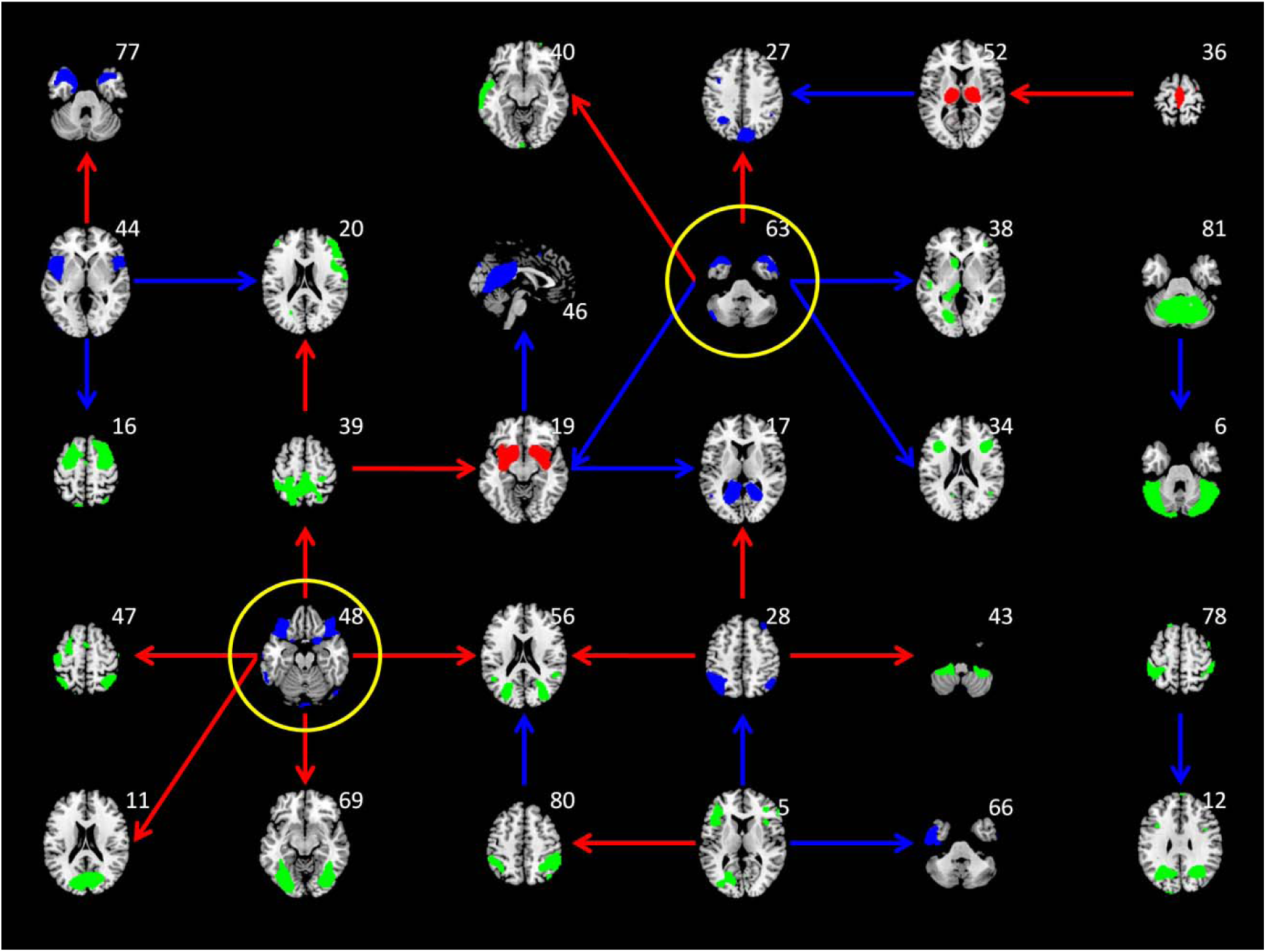
Interregional causal influences network at *λ* of 0.162 using LASSO (least absolute shrinkage and selection operator) regression. The colors of the independent component maps represent increased (red), decreased (blue), and non-significant (green) age effects on metabolic activity. The colors of the arrows represent positive (red) and negative (blue) interregional influences, respectively. The maps highlighted with yellow circle represent hub regions in the network.

To better illustrate the topology of the interregional influencing network and to highlight the regions that are more influencing or influenced to other regions, we plotted the first giant components of the interregional influencing network using force layout at *λ* = 0.162, and also at more liberal thresholds of *λ* = 0.142 and *λ* = 0.122 (Figure 8). The node sizes represent the out-degree or in-degree of a node in the network in the upper and middle panels, respectively. It can be seen that the nodes with large out-degree were in general the regions with decreased metabolic activity (blue nodes). The red arrows highlighted the two nodes that had 5 out-degrees at *λ* = 0.162 and remained among the highest out-degree nodes at the lower *λ* values. These two nodes were also highlighted in Figure 7, which covered the bilateral orbitofrontal cortex (IC# 48) and the bilateral anterior temporal lobe (IC# 63). While in terms of in-degree, there were no clear regions that had exceptionally high in-degree compared with other nodes. The node with high in-degree had no apparent age effects (green nodes) or had increased metabolic activity with age (brown nodes). The distributions of nodal out- and in-degree confirmed that the out-degree distributions had heavy tailed distributions compared with the in-degree distributions (lower panels in Figure 8).

**Figure 8.**
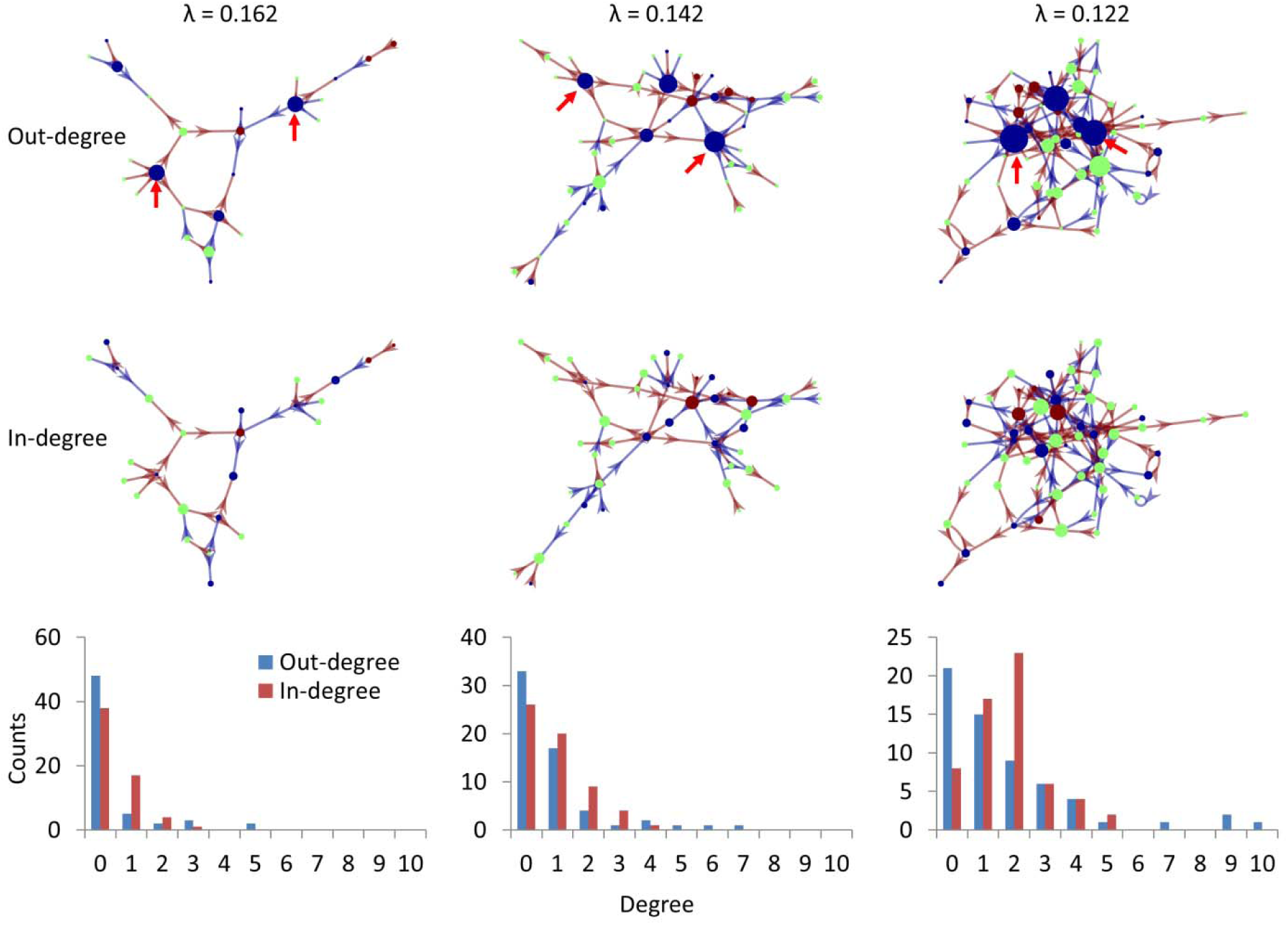
The giant component of the causal interregional influencing network of metabolic activity at different *λ* levels with the sizes of the nodes reflecting out-degree (top row) and in-degree (middle row). The bottom row shows the degree distributions of the whole influencing network at the three *λ* levels. Brown and blue arrows indicate positive and negative influences identified using LASSO regression. Blue, green, and brown regions indicate positive, none, and negative age effects on regional metabolic activity. The red arrows highlight the two influencing nods at different *λ* levels.

## 4. Discussion

By applying autoregressive model on longitudinal FDG-PET data, the current study demonstrated causal interregional influences of metabolic activity during normal aging. Several ICs with significant reduced metabolic activity in aging, including the orbital frontal cortex and anterior temporal lobe, causally influenced many other ICs. In contrast, the influenced ICs were more widespread and with less local aging effects or even with relatively increased metabolic activity. To the best of our knowledge, this is the first study to demonstrate longitudinal interregional causal influences of brain activity during aging at the time scale of a year.

Consistent with our predictions of interregional spreads of age effects, the influencing ICs usually had decreased metabolic activity. On the other hand, the influenced regions were not restricted to the regions with reduced metabolic activity in aging. Indeed, the ICs that had relatively greater in-degree values than other ICs were usually without apparent age effects, or even with relatively increased metabolic activity, e.g. the basal ganglia and thalamus. Therefore, the causal interregional influences in general reflected the spread of age effects from brain regions that had already declined to regions that are declining or relatively preserved. We note that the absence of regional age effects should be interpreted with caution, because the removal of global effects during calculation of regional age effects could have removed significant age effects that were similar to the global effects.

The interpretation of the causal influences need to consider both the sign of the causal influences and the regional age effects. A positive influence indicates that the metabolic activity in region A at the current time point positively predicts the metabolic activity in region B at the next time point. While a negative influence indicates a negative prediction. If the two regions are both decreasing during aging, then a positive influence may indicate a spread of metabolic activity decline between the two regions. On the other hand, if region A decreases but region B shows relatively increased metabolic activity, and there is a negative influence between A and B, then it may indicate a compensation of region B that is resulted from the declined function of region A. A close look at the patterns of the directions of the local and interregional effects indicated that most of the effects observed were consistent with the spatial spread or compensation interpretations. That is, the influences between two decreased regions were all positive, and the influences between one decreased region and one increased regions were all negative.

The current analysis identified several hubs that influenced other brain regions, most prominently the anterior temporal lobe and orbital frontal cortex. The anterior temporal lobe (IC 63) is connected to several major white matter tracts such the cingulum, inferior longitudinal fasciculus, and uncinate fasciculus (Catani & Thiebaut de Schotten, 2008), which could support its influencing role to other regions such as the subcortical regions, inferior frontal cortex, and left temporal cortex. To better characterize its functional correlates, we submitted the IC map into NeuroVault (NeuroVault, RRID:SCR_003806; https://neurovault.org), and decoded the functions of these maps using large-scale meta-analytic data from Neurosynth (NeuroSynth, RRID:SCR_006798; http://neurosynth.org/) (Rubin et al., 2017). The first five functional terms were all about language and semantic processing (See Supplementary Table S2). Studies also showed that electrical stimulation of the anterior temporal lobe can improve proper name recalls in aging (Ross, McCoy, Coslett, Olson, & Wolk, 2011), and bilingualism can protect the integrity of anterior temporal lobe in aging (Abutalebi et al., 2014). Taken together, the results suggest that language process might be an important factor modulating brain aging.

The orbital frontal cortex (IC 48) is connected to the uncinate fasciculus and inferior fronto-occipital fasciculus (Catani & Thiebaut de Schotten, 2008), which could support its influences to the posterior visual regions. The functional words related to the orbitofrontal IC were mainly about emotional processing (Supplementary Table S2). In older population, smaller orbitofrontal volumes are shown to be associated with depression (Lai, Payne, Byrum, Steffens, & Krishnan, 2000; W. D. Taylor et al., 2003). Taken together, emotional process might also be an important factor modulating brain aging. However, although previous studies have shown associations between resting-state brain activity and task activations (Di, Kannurpatti, Rypma, & Biswal, 2013; Yuan et al., 2013), the extent to what resting-state brain activity can reflect certain brain functions are still largely unknown. Further studies might need to design proper tasks to better link functions to brain activations. A limitation of the current analysis is the potential confounding effect due to partial volume (Bonte et al., 2017; Rousset, Ma, & Evans, 1998), i.e. whether an observed effect is due to the changes of *bona fide* metabolic activity or the changes of underlying gray matter volume. However, the following reasons make the partial volume confounding less problematic. First, the current analysis adopted within subject comparison, which has already minimized the partial volume effects due to inter subject anatomical variability. Second, we applied ICA analysis to identify independent sources of metabolic variability. Some components that were likely due to enlargement of ventricle and are spatially overlapped with the included ICs, have been already removed. For example, there was an IC largely located in ventricle area (IC 79 in supplementary Figure S9) but with substantial overlaps with the ICs of the thalamus and basal ganglia (supplementary Figure S5). The IC 79 had the second strongest negative age effect among all the ICs. The included ICs that had spatial overlap with this IC showed no age effects or even positive age effects, suggesting that the partial volume effects associated with enlarged ventricle have been minimized in these ICs. Third, even though the observed causal influences may still somehow contributed by the residual partial volume effects, the causal influences of volumetric reductions may still be important findings for understanding brain aging. The structural MRI images are available in the ADNI dataset, but were not always acquired at the same time point as the PET images, making the incorporation of MRI images in the model difficult. Future studies should certainly consider taking into account of anatomical information in the analysis. Indeed, it may be theoretically more important to study the interaction or causal influences between brain anatomy and functions in aging. According to the compensation model, the reduction of gray matter will lead to elevated functional responses, which then give rise to less affected behavioral performances (Gregory et al., 2018; Reuter-Lorenz & Park, 2014; Shafto & Tyler, 2014). A direct examination of causal influences ainfluences among local and interregional gray matter structures, functions, and behavioral performances may provide more insight to the dynamic of compensation process in aging.

One strength of the current analysis approach is that we adopted multivariate methods and LASSO to include all the ICs in the predicting models, which in theory can prevent identifying ICs that have indirect predicting effects to the target (Smith et al., 2011; Tang et al., 2012). On the other hand, there are also several simplifications of the Granger causality analysis, such as the inclusion of only the first order model and the assumption of equal time steps. Since the current study is the first to explore the causality in the aging process, the time lag of the aging progression is still largely unknown and bear further studies. But practically due to the limited availability of longitudinal data, this question is difficult to solve at the current stage. Regarding the variable time steps of the time series, we added the intersession interval as a covariate to minimize the effects, which is similar to a previous work (Jiang et al., 2018). More sophisticated models, such as generative model and differential equation based method (G. Ziegler, Penny, Ridgway, Ourselin, & Friston, 2015; Gabriel Ziegler, Ridgway, Blakemore, Ashburner, & Penny, 2017), may be used in future to better characterize the causal effects.

## 5. Conclusion

By applying Granger causality analysis on longitudinal FDG-PET images of healthy old participants at a time step of one year, the current analysis revealed interregional causal influences during aging. Several regions with reductions in local metabolic activity during aging, including the bilateral anterior temporal lobe and orbitofrontal cortex, showed causal influences to other regions, supporting an interregional spread of age effects in the brain. The current analysis and results could add new insights to the neurocognitive aging literature about interregional interactions during the aging process.

## Acknowledgements

Data collection and sharing for this project was funded by the Alzheimer’s Disease Neuroimaging Initiative (ADNI) (National Institutes of Health Grant U01 AG024904) and DOD ADNI (Department of Defense award number W81XWH-12-2-0012). ADNI is funded by the National Institute on Aging, the National Institute of Biomedical Imaging and Bioengineering, and through generous contributions from the following: AbbVie, Alzheimer’s Association; Alzheimer’s Drug Discovery Foundation; Araclon Biotech; BioClinica, Inc.; Biogen; Bristol-Myers Squibb Company; CereSpir, Inc.; Cogstate; Eisai Inc.; Elan Pharmaceuticals, Inc.; Eli Lilly and Company; EuroImmun; F. Hoffmann-La Roche Ltd and its affiliated company Genentech, Inc.; Fujirebio; GE Healthcare; IXICO Ltd.; Janssen Alzheimer Immunotherapy Research & Development, LLC.; Johnson & Johnson Pharmaceutical Research & Development LLC.; Lumosity; Lundbeck; Merck & Co., Inc.; Meso Scale Diagnostics, LLC.; NeuroRx Research; Neurotrack Technologies; Novartis Pharmaceuticals Corporation; Pfizer Inc.; Piramal Imaging; Servier; Takeda Pharmaceutical Company; and Transition Therapeutics. The Canadian Institutes of Health Research is providing funds to support ADNI clinical sites in Canada. Private sector contributions are facilitated by the Foundation for the National Institutes of Health (www.fnih.org). The grantee organization is the Northern California Institute for Research and Education, and the study is coordinated by the Alzheimer’s Therapeutic Research Institute at the University of Southern California. ADNI data are disseminated by the Laboratory for Neuro Imaging at the University of Southern California.

## Author contributions

X.D. conceived the idea, developed and performed the data analysis. All authors discussed the results, and contributed to the final manuscript.

## Conflict of interest statement

The authors declare that there is no conflict of interest regarding the publication of this article.

## Supplemental materials

Supplemental materials include one word file containing nine supplementary Figures and two supplementary Tables. Other supporting information can be found at https://osf.io/4a3vt/.

## Notes

https://osf.io/4a3vt/

