## Supplemental Materials for "Interregional causal influences of brain metabolic activity reveal the spread of aging effects during normal aging"

This document contains supporting information for the manuscript titled “Interregional causal influences of brain metabolic activity reveal the spread of aging effects in normal aging”. It includes:

Figure S1 – S8: Independent component (IC) maps of metabolic activity of the eight domains.

Figure S9: An independent component (IC) map of metabolic activity of an example noise component.

Table S1: List of all the independent components (ICs) used in the analysis and their mainly covered brain structures.

Table S2: Neurosynth decoding results of the four hub ICs identified in the causality analysis.

More information can be found at: <https://osf.io/4a3vt/?view_only=74b5ca53938c4dda9eb51ee338f9a0ab>

**Figure S1** Independent component (IC) maps of metabolic activity of the cerebellum. The maps in red, blue, and green represent positive, negative, and no age effects on local metabolic activity. The maps were thresholded at beta > 2.3. The numbers at the bottom represent z coordinates in Montreal Neurological Institute (MNI) space.

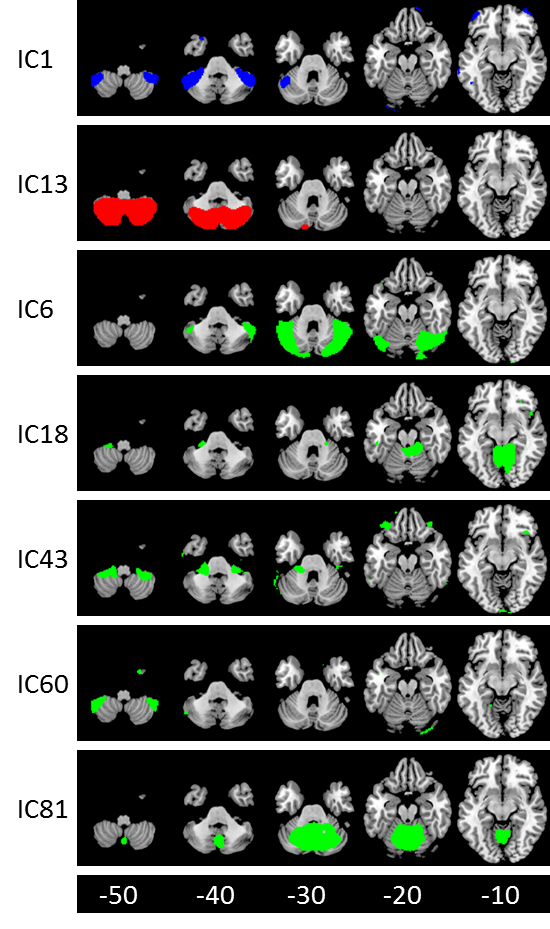

**Figure S2** Independent component (IC) maps of metabolic activity of the visual region. The maps in red, blue, and green represent positive, negative, and no age effects on local metabolic activity. The maps were thresholded at beta > 2.3. The numbers at the bottom represent z coordinates in Montreal Neurological Institute (MNI) space.

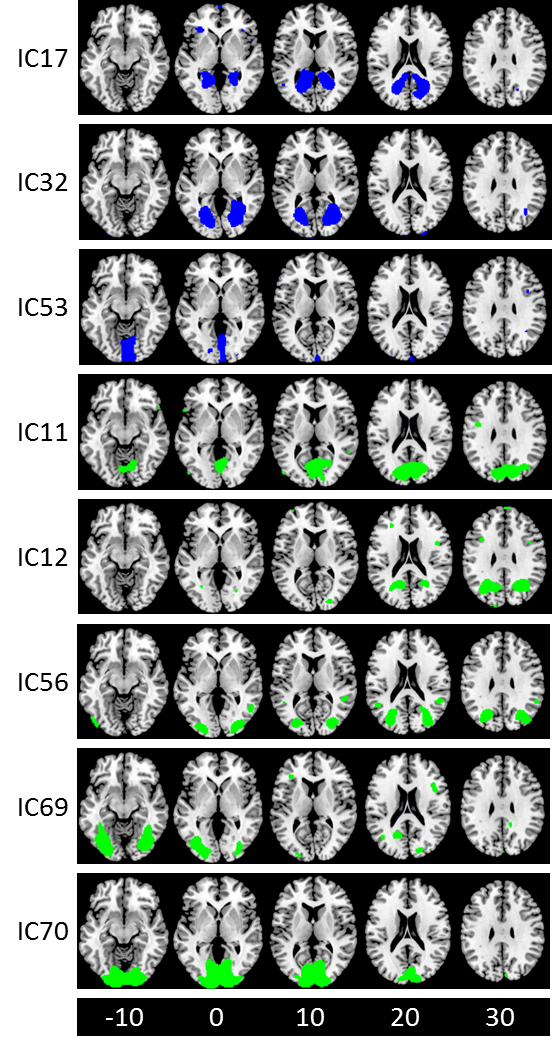

**Figure S3** Independent component (IC) maps of metabolic activity of the parietal region. The maps in red, blue, and green represent positive, negative, and no age effects on local metabolic activity. The maps were thresholded at beta > 2.3. The numbers at the bottom represent z coordinates in Montreal Neurological Institute (MNI) space.

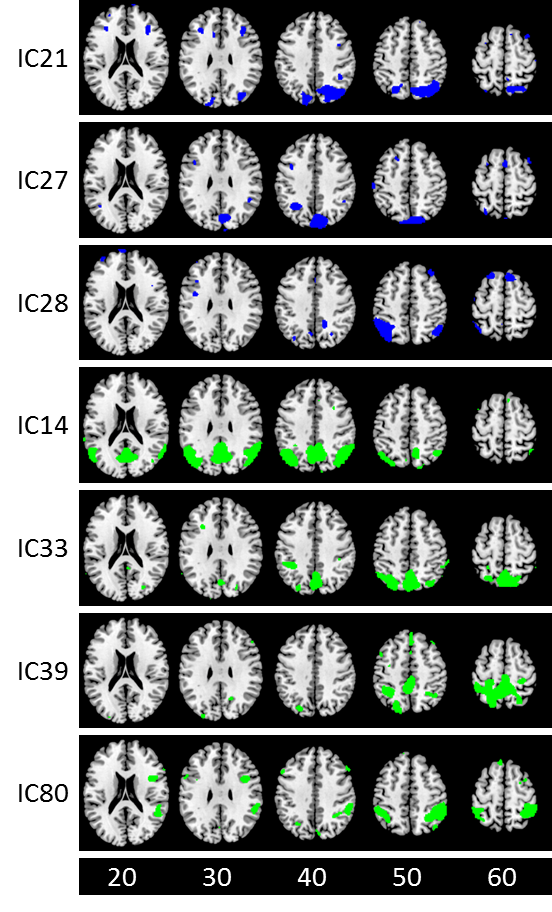

**Figure S4** Independent component (IC) maps of metabolic activity of the temporal region. The maps in red, blue, and green represent positive, negative, and no age effects on local metabolic activity. The maps were thresholded at beta > 2.3. The numbers at the bottom represent z coordinates in Montreal Neurological Institute (MNI) space.

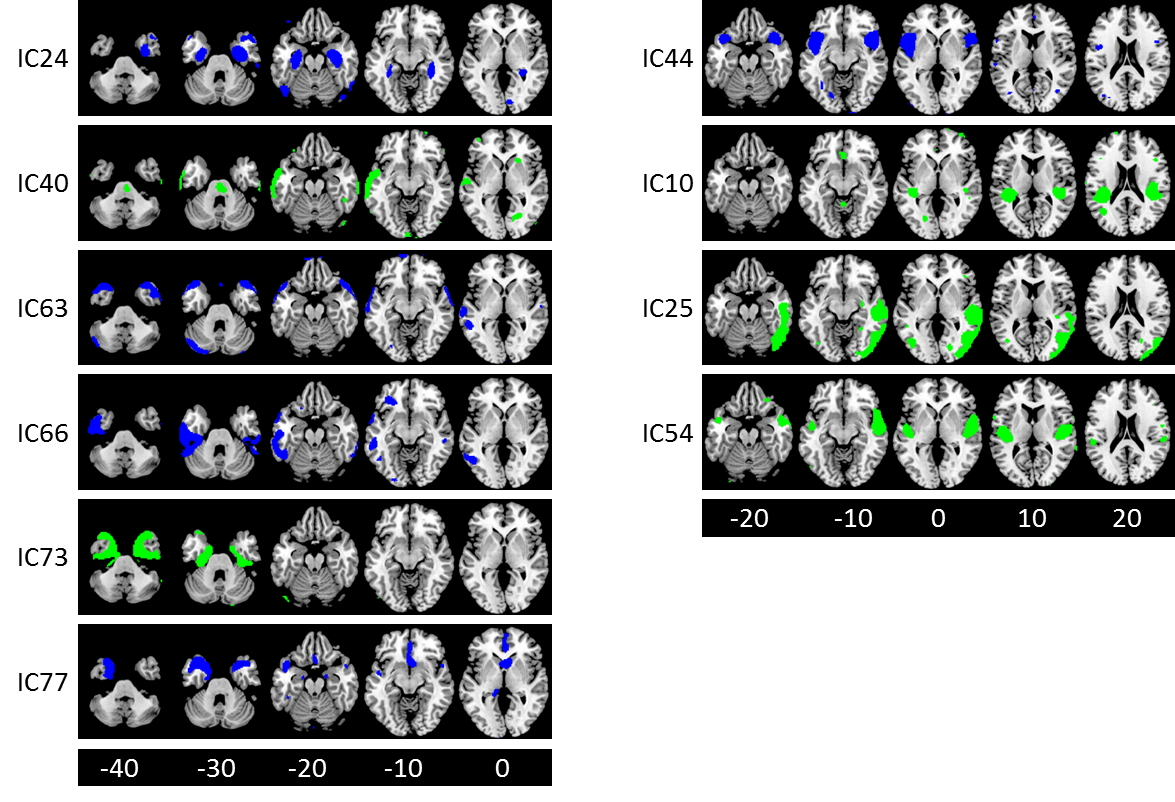

**Figure S5** Independent component (IC) maps of metabolic activity of the subcortical regions. The maps in red, blue, and green represent positive, negative, and no age effects on local metabolic activity. The maps were thresholded at beta > 2.3. The numbers at the bottom represent z coordinates in Montreal Neurological Institute (MNI) space.

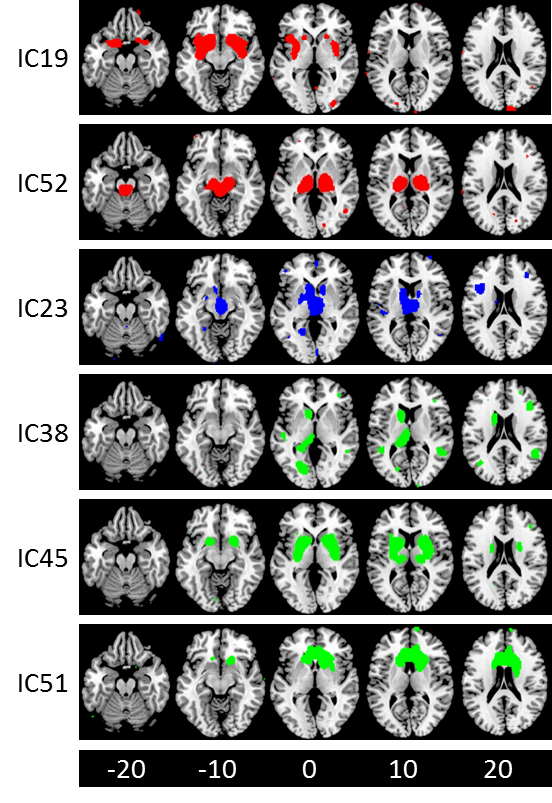

**Figure S6** Independent component (IC) maps of metabolic activity of the frontal region. The maps in red, blue, and green represent positive, negative, and no age effects on local metabolic activity. The maps were thresholded at beta > 2.3. The numbers at the bottom represent z coordinates in Montreal Neurological Institute (MNI) space.

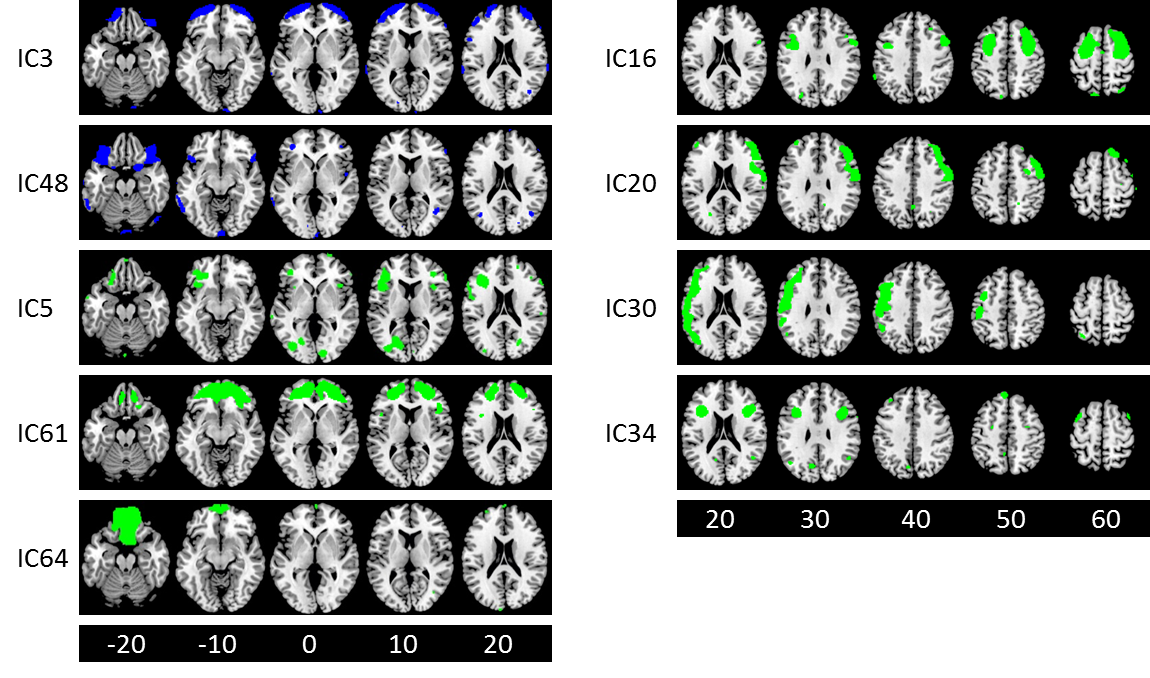

**Figure S7** Independent component (IC) maps of metabolic activity of the sensorimotor regions. The maps in red, blue, and green represent positive, negative, and no age effects on local metabolic activity. The maps were thresholded at beta > 2.3. The numbers at the bottom represent z coordinates in Montreal Neurological Institute (MNI) space.

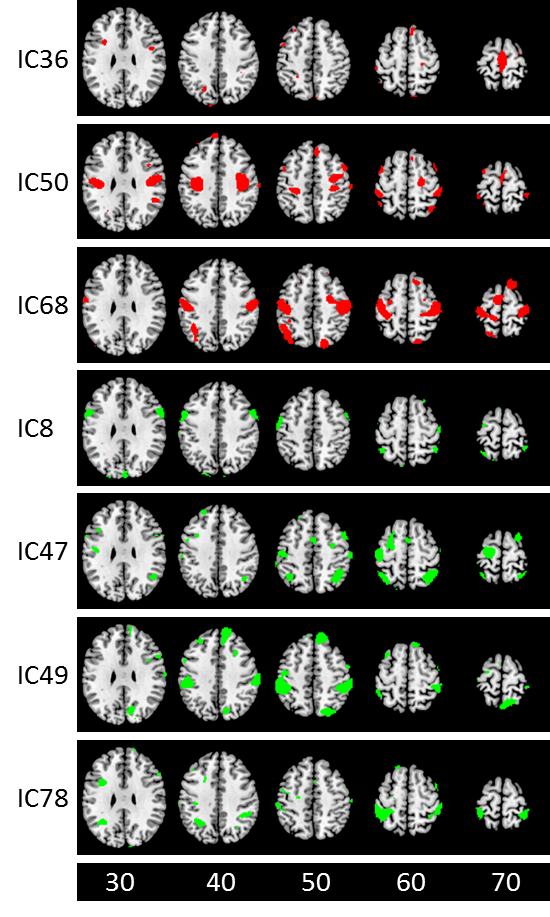

**Figure S8** Independent component (IC) maps of metabolic activity of the cingulate region. The maps in red, blue, and green represent positive, negative, and no age effects on local metabolic activity. The maps were thresholded at beta > 2.3. The numbers at the bottom represent z coordinates in Montreal Neurological Institute (MNI) space. The right panels represent the medial axial slice (x = 0 in MNI space) of each IC.

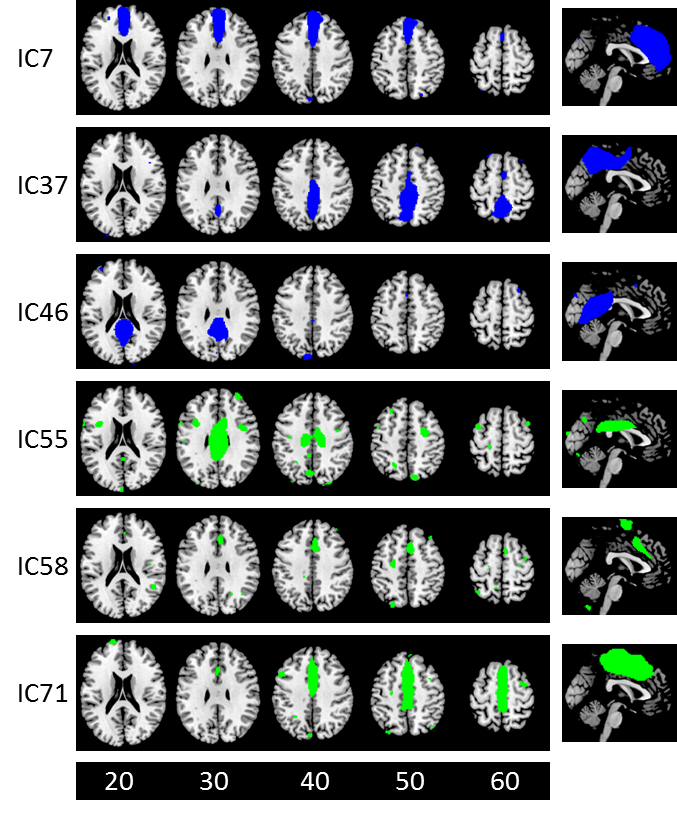

**Figure S9** An independent component (IC) map of metabolic activity of an example noise component, which mainly covers the ventricle. This map spatially overlaps with several subcortical ICs shown in Figure S5. The maps were thresholded at beta > 2.3. The numbers at the bottom represent z coordinates in Montreal Neurological Institute (MNI) space.

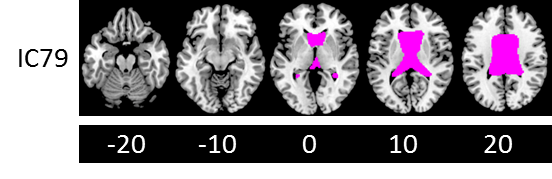

**Table S1** List of independent components (ICs) for the eight domains and their mainly covered brain structures.

| IC # | Label | Domain |
| --- | --- | --- |
| 1 | Lobule VII and VIII, hemisphere | Cerebellum |
| 6 | Lobule VII crus | Cerebellum |
| 13 | Lobule VII, hemisphere and crus | Cerebellum |
| 18 | Vermis | Cerebellum |
| 43 | Lobule VII and VIII, hemisphere | Cerebellum |
| 60 | Lobule VII and VIII, hemisphere | Cerebellum |
| 81 | Lobule VII crus, Lobule VI hemisphere, vermis | Cerebellum |
| 11 | Lingual gyrus, calcarine sulcus, cuneus | Visual |
| 12 | Superior and middle occipital gyrus | Visual |
| 17 | Lingual gyrus, calcarine sulcus, cuneus | Visual |
| 32 | Lingual gyrus, calcarine sulcus | Visual |
| 53 | Lingual gyrus | Visual |
| 56 | Superior and middle occipital gyrus | Visual |
| 69 | Inferior occipital and fusiform gyrus | Visual |
| 70 | Lingual gyrus, calcarine sulcus, cuneus | Visual |
| 14 | Angular gyrus, posterior cingulate cortex | Parietal |
| 21 | Superior parietal lobule, angular gyrus | Parietal |
| 27 | Precuneus | Parietal |
| 28 | Inferior parietal lobule, precuneus | Parietal |
| 33 | Superior parietal lobule, precuneus | Parietal |
| 39 | Superior parietal lobule, precuneus | Parietal |
| 80 | Inferior parietal lobule | Parietal |
| 10 | Rolandic operculum | Temporal |
| 24 | Hipocampal, parahipocampal, and furisom gyrus | Temporal |
| 25 | Right temporal and occipital lobe | Temporal |
| 40 | Left middle temporal gyrus | Temporal |
| 44 | Insula, inferior frontal gyrus, superior temporal pole | Temporal |
| 54 | Superior temporal gyrus | Temporal |
| 63 | Temporal pole | Temporal |
| 66 | Left inferior temporal gyrus | Temporal |
| 73 | Temporal pole, parahipocampal gyrus | Temporal |
| 77 | Temporal pole, medial orbitofrontal cortex | Temporal |
| 19 | Basal ganglia, amygdala, insula | Subcortical |
| 23 | Thalamus, caudate | Subcortical |
| 38 | Left thalamus, caudate, lingual gyrus | Subcortical |
| 45 | Thalamus, basal ganlia | Subcortical |
| 51 | Caudate, anterior cingulate cortex | Subcortical |
| 52 | Thalamus, brainstem | Subcortical |
| 3 | Lateral frontal pole | Frontal |
| 5 | Left inferior frontal gyrus, left middle occipital gyrus | Frontal |
| 16 | Superior and middle frontal gyrus | Frontal |
| 20 | Right inferior and middle frontal gyrus | Frontal |
| 30 | Left inferior and middle frontal gyrus extending to temporal lobe | Frontal |
| 34 | Inferior frontal gyrus | Frontal |
| 48 | Orbitofrontal cortex, superior temporal pole | Frontal |
| 61 | Medial frontal pole | Frontal |
| 64 | Ventromedial prefrontal cortex | Frontal |
| 8 | Precentral gyrus | Motor |
| 36 | Supplementary motor area, paracentral lobule | Motor |
| 47 | Postcentral gyrus, inferior parietal lobule | Motor |
| 49 | Postcentral gyrus, inferior parietal lobule | Motor |
| 50 | Postcentral and precentral gyrus | Motor |
| 68 | Postcentral gyrus, angular gyrus | Motor |
| 78 | Postcentral gyrus | Motor |
| 7 | Anterior cingulate cortex, medial frontal gyrus | Cingular |
| 37 | Posterior and middle cingulate cortex, precuneus | Cingular |
| 46 | Posterior cingulate cortex, precuneus, lingual gyrus | Cingular |
| 55 | Middle and posterior cingulate cortex | Cingular |
| 58 | Middle cingulate cortex, supplementary motor area | Cingular |
| 71 | Middle cingulate cortex, supplementary motor area | Cingular |

**Table S2** Top five functional keywords of the two influencing hub IC maps decoded from Neurosynth.

| ID | 48 | 63 |
| --- | --- | --- |
| Label | Orbital frontal cortex | Anterior temporal lobe |
| Keywords | Dementia | Voice |
|  | Prospective | Speaker |
|  | Emotional | Spoken |
|  | Anger | Sentences |
|  | Cognitive emotional | Comprehension |
